# Predicting structures of large protein assemblies using combinatorial assembly algorithm and AlphaFold2

**DOI:** 10.1101/2023.05.16.541003

**Authors:** Ben Shor, Dina Schneidman-Duhovny

## Abstract

Deep learning models, such as AlphaFold2 and RosettaFold, enable high-accuracy protein structure prediction. However, large protein complexes are still challenging to predict due to their size and the complexity of interactions between multiple subunits. Here we present CombFold, a combinatorial and hierarchical assembly algorithm for predicting structures of large protein complexes utilizing pairwise interactions between subunits predicted by AlphaFold2. CombFold accurately predicted (TM-score > 0.7) 72% of the complexes among the Top-10 predictions in two datasets of 60 large, asymmetric assemblies. Moreover, the structural coverage of predicted complexes was 20% higher compared to corresponding PDB entries. We applied the method on complexes from Complex Portal with known stoichiometry but without known structure and obtained high-confidence predictions. CombFold supports the integration of distance restraints based on crosslinking mass spectrometry and fast enumeration of possible complex stoichiometries. CombFold’s high accuracy makes it a promising tool for expanding structural coverage beyond monomeric proteins.

## Introduction

Most proteins function as multimolecular assemblies in the cells. There are on average a few dozen interactions per protein^1–3^. These assemblies perform important functions, such as energy transduction^4^, transport^5^, and signal transduction^6^. The determination of the 3D structures of these assemblies is critical for understanding their function and evolution, interpreting the effects of mutations, and potential applications in drug discovery. The large size of some assemblies and conformational heterogeneity poses challenges for traditional structural characterization techniques, such as X-ray crystallography and NMR spectroscopy. While progress has been made using cryo-Electron Microscopy (cryo-EM), high-throughput structure determination of large assemblies is still challenging.

Recently deep learning techniques significantly advanced our ability to predict high-accuracy protein structures. One of the most significant advancements was the release of AlphaFold2^7^ and RosettaFold^8^. While AlphaFold2 was designed to predict single-chain proteins, it can also apply to predict protein complexes using the same architecture. Soon after its release, several techniques were developed to use AlphaFold2 to predict multi-chain protein complexes - first by using a linker^9^ and later by offsetting the residue index^10^. Similar techniques were used for the training of AlphaFold-Multimer (AFM)^11^ which is able to predict multimeric complexes with high accuracy using paired and padded multiple sequence alignment (MSA). On several pairwise protein-protein docking benchmarks AFM achieves a success rate of 40-70% for complexes consisting of two to nine chains up to 1,536 in total length^11–13^.

However, AFM application for predicting structures of large assemblies is still challenging^12, 13^. The first difficulty is the requirement for significant resources, such as a graphical processing unit (GPU) with a large memory size. Currently, common GPUs have no more than 20 gigabytes of memory, enabling the prediction of complexes up to 1,800 and 3,000 amino acids for AFMv2 and AFMv3, respectively. Also, as AFM memory usage increases roughly quadratically with the number of amino acids^7^, any potential hardware advancements in the future are unlikely to have a significant impact. This limits the practical capability of many researchers to predict structures of large size, leaving many macromolecular complexes without a structure prediction. The second difficulty is sampling with a large number of restraints: as the number of chains and amino acids increases, the number of residue-residue contacts and distance restraints to optimize increases as well, making it harder for the model to converge to accurate structures. Large, multimolecular complex prediction is an out-of-domain inference setup for AFM since it was trained only on cropped regions and thus is not expected to perform well. The third difficulty is that AFM converges to a single (sometimes incorrect) structure (for each of the five available trained models) and it is highly challenging to obtain a diverse set of predictions for the same target^14^.

Prior to the deep learning revolution, methods developed for the assembly of multi-protein complexes could be divided into two main categories. The first category is integrative modeling methods that mainly rely on experimental data^15, 16^, and the second is docking-based methods that rely on pairwise protein-protein docking^17–19^. Integrative modeling methods rely on information from multiple sources, such as crosslinking mass spectrometry, Förster resonance energy transfer, co-evolution, cryo-EM, and small angle x-ray scattering to compute models. This information is converted into spatial restraints and combined into an integrative modeling approach^20, 21^, using specialized software packages^15, 22, 23^ to generate a set of structural models that are consistent with it. The generation of candidate models is often performed by a Monte Carlo search of the conformational space. Docking-based methods significantly rely on pairwise protein-protein docking for the prediction of complexes^24–26^ and do not require additional input information. In pairwise docking, the two input proteins are docked to one another using geometric shape and physicochemical complementarity. The main problem is that they sample thousands of docked configurations. While the correct ones are usually sampled, it is difficult to rank them as top-scoring. Typically, pairwise docking methods succeed in ranking a correct model among the Top-10 best scoring in 25-40% of the cases^27, 28^. This low accuracy further complicates the multi-protein assembly stage, where methods have to consider a large number of pairwise protein-protein docking models. For example, Multi-LZerD^18^ builds the multimolecular assembly by applying a stochastic search driven by a genetic algorithm. Kuzu et al.^19^ construct the multimolecular complex iteratively, where a single subunit is added to the subassembly in each iteration. The CombDock method is hierarchical and combinatorial^17, 29^. The complexes are constructed hierarchically by generating subassemblies of two or more subunits. At each stage, subassemblies are connected using pairwise docking configurations between subunits. Due to multiple possible hierarchical assembly pathways, the algorithm combinatorially enumerates assembly trees. Since the algorithms used for docking and scoring pairwise interactions have low accuracy, it is difficult to reach high accuracy in multi-subunit docking.

The recently developed MoLPC method relies on AlphaFold2 to produce configurations for pairs and triplets of chains and assemble them using Monte Carlo Tree Search^30^. However, the approach is applicable mainly to homomeric complexes with a success rate of ∼30%. Inspired by this work, here we combine AlphaFold2 with a deterministic combinatorial assembly algorithm^17, 29^. Our new method, CombFold, uses a small number of pairwise subunit interactions generated by AlphaFold2 for assembly instead of thousands generated by docking. The hierarchical and combinatorial assembly stage exhaustively enumerates possible assembly trees, maximizing the probability of correctly assembling the complex based on pairwise AlphaFold2 interactions. We validate our approach on two benchmarks of large heteromeric assemblies (up to 30 chains and 18,000 amino acids) and obtain a Top-1 success rate of 62% and Top-10 success rate of 72% (TM-score > 0.7). Moreover, CombFold is able to increase the structural coverage by 20% relative to experimental structures in our benchmarks. We also test the method on the benchmark of homomeric complexes used for MoLPC validation and obtain a Top-1 success rate of 57%. CombFold successfully assembles six out of seven CASP15 targets with over 3,000 amino acids. We apply the method on a set of complexes with known stoichiometry and without known structure from Complex Portal^31^ and obtain confident predictions.

## Results

### Methods summary

The input to CombFold is the subunit sequences and optionally distance restraints, the output is a set of assembled structures. A subunit can be a single chain or a domain. The approach is based on combinatorial and hierarchical assembly via pairwise interactions. In principle, there is no limitation on complex size, as the complex can be divided into subunits suited for the GPU memory limit, and our current implementation supports up to 64 subunits. CombFold works in three major stages: (i) generation of pairwise subunit interactions by AFM, (ii) creation of a unified representation of subunits and interactions, and (iii) combinatorial assembly of subunits (Fig. 1).

**Figure 1.**
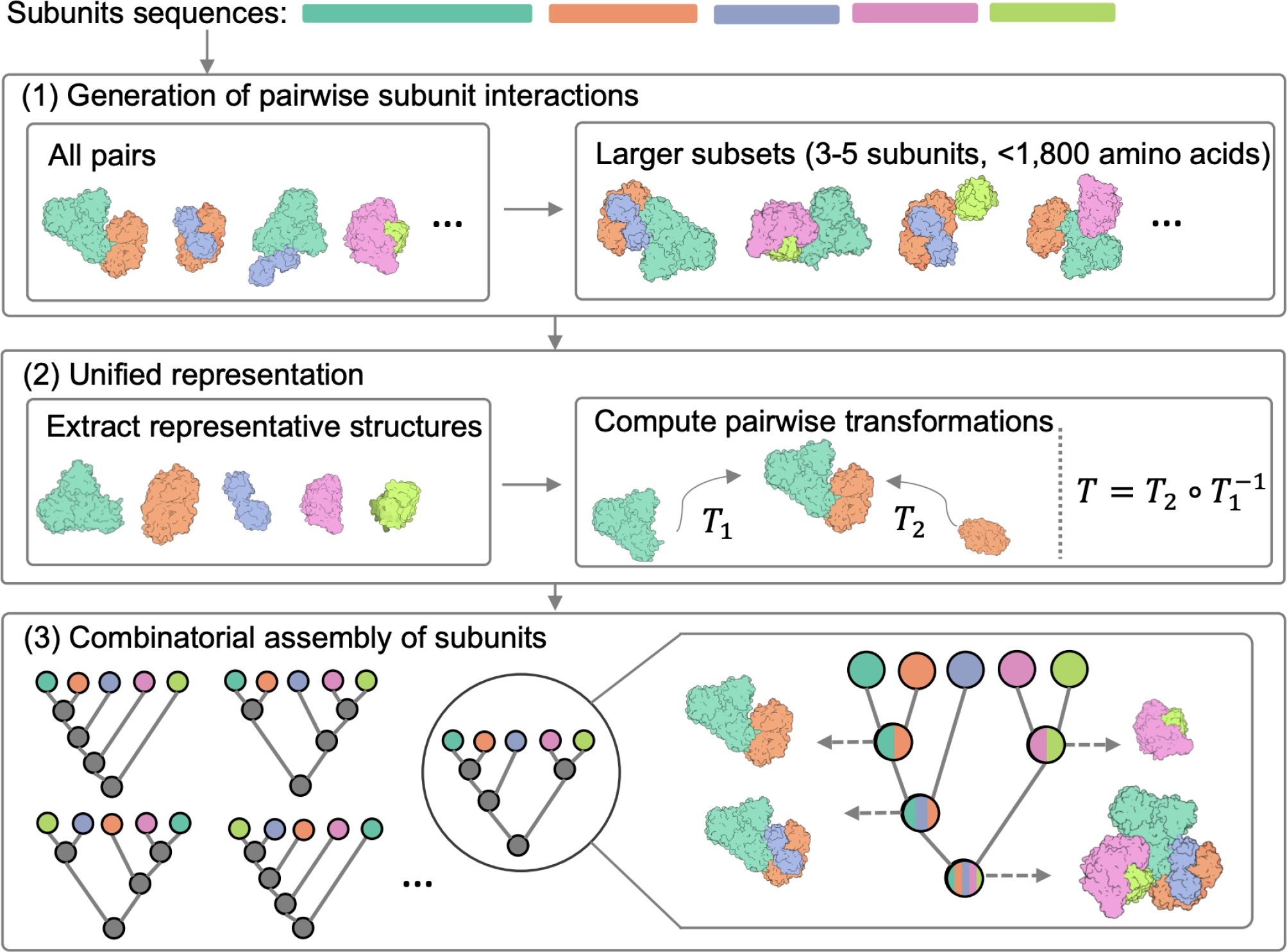
The three stages of the CombFold assembly algorithm. The input is the sequences of the subunits in the complex. (1) Structure prediction of all pairwise and some larger subunit subsets using AFM. (2) Selection of representative subunit structures out of all predicted structures, followed by computation of all pairwise transformations present in predicted structures relative to the representative structures. (3) Combinatorial and hierarchical assembly of subunit structures using the computed pairwise transformations. In each iteration, new subcomplexes are assembled using a pairwise transformation to join two previously created subcomplexes.

In the first stage, we apply AFM to all possible subunit pairings. Following this, we create three additional AFM models for each subunit, ranging in size from three to five subunits, that include subunits with which the given subunit had the highest confidence-scored predicted pairwise interactions (Methods). The underlying concept is that some groups of more than two subunits form intertwined structures and therefore all of them should be predicted as a single model by AFM (Methods).

In the second stage, to prepare input for the third assembly stage, a single representative structure for each subunit is selected and the transformations between representative subunits are calculated. This is required since there are multiple AFM structures for each subunit from pairwise AFM runs and their enumeration during the assembly stage is intractable. The representative subunit structures are extracted from the predicted modeled subcomplexes according to the maximal average plDDT score for this subunit. Next, we use all interacting subunit pairs (Cα-Cα distance < 8Å) from AFM models to extract pairwise transformations (rotation and translation in 3D) between their representative structures in the global reference frame. The representation of the input by representative subunit structures and transformations between them enables us to apply the combinatorial assembly algorithm with AFM interactions instead of docking-based ones. Each transformation is coupled with a score based on AFM’s PAE score (Methods).

In the third stage, we use *N* representative subunit structures, the pairwise transformations between them, and optionally distance restraints, for the hierarchical and combinatorial assembly of the entire complex. Distance restraints can originate from crosslinking mass spectrometry, Förster resonance energy transfer, or other sources of information^32–35^. If a protein chain is divided into subunits (eg. domains), distance constraints are added to enforce sequence connectivity. This combinatorial assembly stage consists of *N* iterations, where in the *i*-th iteration we construct *K* subcomplexes of size *i*. The value of *K* has to be large enough to contain a variety of subcomplexes. Subcomplexes of size *i* are constructed from pairs of previously computed subcomplexes of size 1 to *i-1*. For example, a subcomplex of size *i* can be computed by merging subcomplexes of size *3* and *i-3*. We attempt to merge a pair of subcomplexes if they do not have any shared subunit and the joint number of subunits is *i*. During the merge, new subcomplexes are generated by iterating all subunit pairs (one from each subcomplex) and applying known transformations between those two subunits on the entire subcomplexes. Next, we discard generated subcomplexes with significant steric clashes or chain connectivity violations. Distance restraints satisfaction is calculated and low-scoring subcomplexes are also discarded. The remaining subcomplexes are clustered and scored based on the score of transformations that were used, and the top *K* subcomplexes are saved for the next iterations.

The model confidence score produced by our method is based on the AFM PAE score. Each pairwise interaction (represented by a transformation) has a PAE-based score (Methods). The confidence of an assembled structure is a weighted score of the transformations that were used for assembly, where the weight is proportional to the sizes of the subunit subsets that were merged by each transformation.

### Benchmark datasets

We tested the method on four benchmark datasets (Table 1). We generated a Benchmark 1 dataset aimed to test the method on large heteromeric complexes. Structures with many unique chains usually do not contain significant symmetry which makes them more challenging for assembly, since many different pairwise interactions need to be found and combined. Benchmark 1 contains 35 structures with 5 to 20 chains and at least 5 unique chains per complex, consisting of 1,300 to 8,000 amino acids (Fig. S1a). This dataset includes only complexes released after April 2018, which AFMv2 was not trained on. Benchmark 2 dataset was generated similarly to Benchmark 1 to test the recently released AFMv3. It contains 25 complexes with 5 to 30 chains and 2,000 to 18,000 amino acids (Fig. S1b) that were not in the training set of AFMv3 (released after September 2021). Benchmark 3 dataset was used for benchmarking the MoLPC approach^30^. It contains 153 complexes ranging between 500 and 10,000 amino acids with 10 to 30 chains per complex. This dataset contains mainly symmetric homomers (98 complexes consisting of one unique chain and 27 consisting of two unique chains). Finally, Benchmark 4 dataset contains seven CASP15 targets with more than 3,000 amino acids.

**Table 1:** CombFold evaluation Benchmarks. *For CASP15 targets the fully automated CombFold had a success rate of 57%. Slight modifications, such as dividing chains into domains, increased the success rate to 86%. **We compared CombFold to CASP15 submissions of the Elofsson group that used AFM and MoLPC.

| Benchmark | Target type | # of complexes | # of chains | # of amino acids | Top-1 success rate of CombFold | Top-10 success rate of CombFold | Top-1 success rate of AFM or MoLPC |
| --- | --- | --- | --- | --- | --- | --- | --- |
| 1 | Asymmetric complexes (released after AFMv2 training) | 35 | 5-20 | 1,300-8,000 | 60% | 74% | 26% (AFMv2) |
| 2 | Asymmetric complexes (released after AFMv3 training) | 25 | 5-30 | 2,000-18,000 | 64% | 68% | 36% (AFMv3) |
| 3 | Mostly homomers and symmetric complexes | 153 | 10-30 | 600-10,000 | 57% | 58% | 28% (MoLPC) |
| 4 | CASP15 targets (> 3,000 amino acids) | 7 | 1-27 | 3,000-8,000 | 57%-86%<sup>*</sup> | 57%-86% | 43% (AFM, MoLPC<sup>**</sup>) |

### Accuracy assessment

To evaluate the accuracy of the modeled structures we rely on the TM-score^36^ which assesses the global accuracy of the complex, similar to CASP and MoLPC^30^. Similarly to CAPRI assessment^37^, a model is considered Acceptable-quality if the TM-score is above 0.7 and High-quality if the TM-score is above 0.8. The success rate is measured as a fraction of the benchmark complexes with Acceptable- or High-quality models among the Top-N best-scoring predictions.

### Accuracy on Benchmark 1 (heteromers)

We obtain a Top-1 success rate of 60% for CombFold on this benchmark, accurately modeling 21 out of 35 complexes (Fig. 2a) with TM-score > 0.7. High-quality Top-1 models are produced for 14 complexes (40%). When considering the Top-10 models, the success rate is 74%. Importantly, the predicted confidence correlates with the TM-score (Pearson r = 0.57, Fig. 2b), indicating that it can be used to estimate model accuracy. To determine to which extent the success rate depends on the ability of AFM to produce accurate models for pairwise interactions, we calculate the pairwise connectivity (Methods). As expected, the pairwise connectivity correlates with the TM-score (Pearson r = 0.48, Fig. 2c).

**Figure 2.**
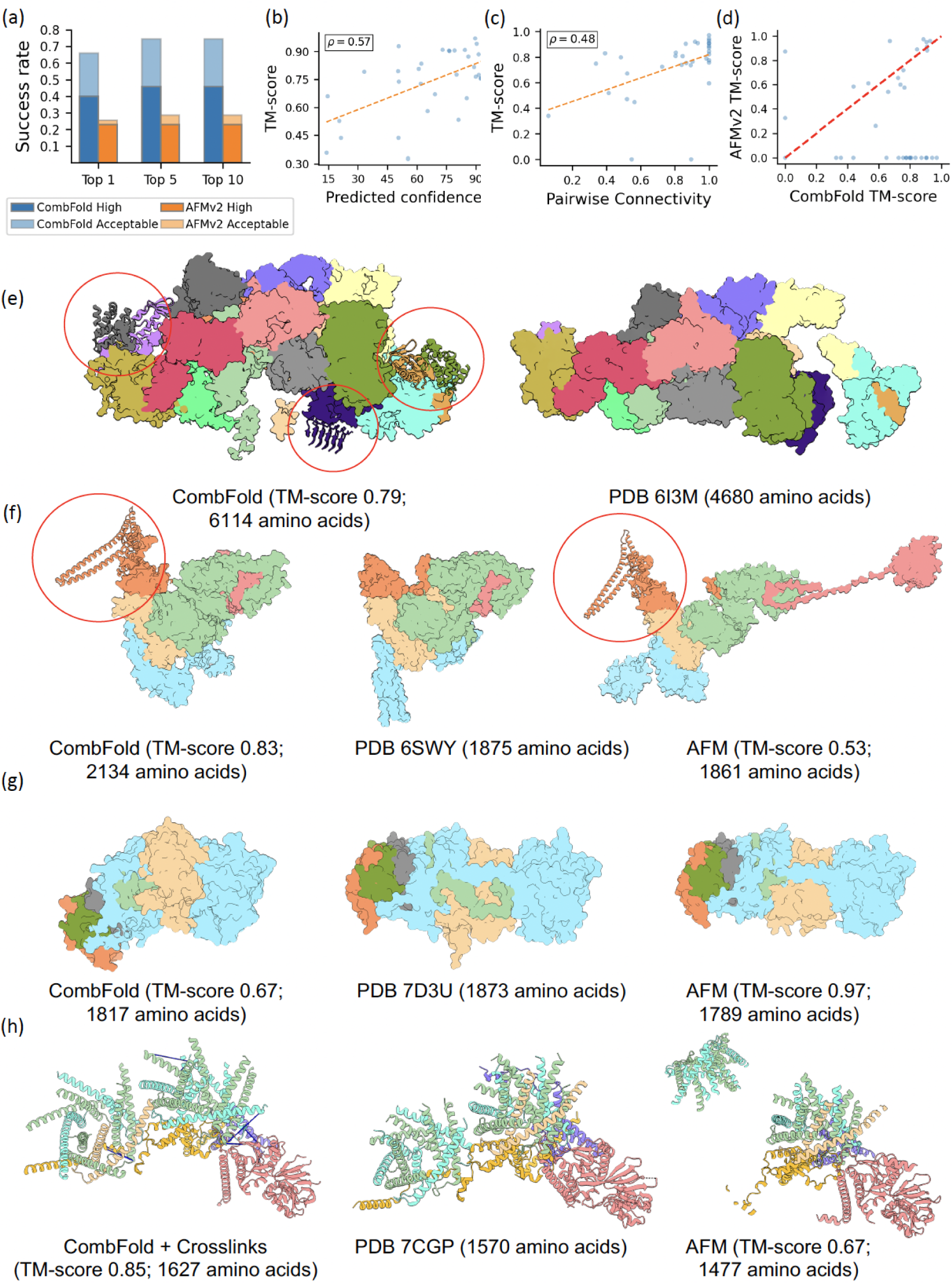
Accuracy of CombFold on Benchmark 1. **(a)** The Top-N (N=1, 5, 10) success rate of CombFold (blue) and AFM (orange). AFM only produces 5 predictions. **(b)** Predicted confidence vs. the TM-score for CombFold. **(c)** Success rate of AFM in producing pairwise interactions as measured by the pairwise connectivity vs. the TM-score of the models produced by CombFold. **(d)** TM-score of AFM models vs. CombFold models. **(e)** eIF2B:eIF2 complex: CombFold model (left) and cryo-EM structure (right). The model contains over 1,500 additional amino acids (marked with red circles). **(f)** GID E3 ubiquitin ligase complex: High-quality CombFold model (left), cryo-EM structure (middle), and inaccurate AFM model (right). **(g)** Multiple resistance and pH adaptation (Mrp) complex: inaccurate CombFold model (left), cryo-EM structure (middle), and High-quality AFM model (right). **(h)** Human mitochondrial translocase TIM22: High-quality model by CombFold, integrating experimental crosslinking data (left), cryo-EM structure (middle), and inaccurate AFM model (right). Crosslinks are shown as blue lines.

We compare CombFold to an end-to-end AFM on all the Benchmark 1 complexes using the A100 GPU card with 40-gigabyte memory. AFM succeeded in producing at least one result for 17 out of 35 complexes with up to 3,700 amino acids. Of these, 10 complexes were modeled with Acceptable- or High-quality, resulting in success rates of 26% and 29% for Top-1 and Top-5 results respectively (Fig. 2a, d).

The largest complex assembled by CombFold was eIF2B:eIF2 (PDB 6I3M, Fig. 2e) which could not be assembled directly with AFM. The CombFold model contains a structural coverage for 6,114 amino acids with plDDT above 50 out of a total of 7,486. In comparison, the experimental cryo-EM structure covers only 4,680 amino acids. The addition of over 1,500 amino acids contains six well-folded domains. This example demonstrates the ability of CombFold to complete unresolved fragments in experimental structures. On average, each assembled complex in this Benchmark contained 20% more amino acids compared to the corresponding PDB entry. GID E3 ubiquitin ligase complex is another example where an additional domain is missing in the experimental structure (PDB 6SWY, Fig. 2f) and is predicted by CombFold with high plDDT. The complex is assembled with a TM-score of 0.83 compared to AFM which produces a model with a TM-score of 0.53. In contrast, the Multiple resistance and pH adaptation (Mrp) complex (PDB 7D3U, Fig. 2g) is assembled with higher accuracy by AFM (TM-score 0.97 vs. 0.67 for CombFold). This is due to the fact that the orientation between the two domains in the largest subunit was not accurately predicted in the representative structure chosen for assembly (Fig. 2g, light blue).

### Accuracy on Benchmark 2 (heteromers)

This benchmark was generated to test CombFold against the recently released AFMv3. We also used AFMv3 to predict the pairwise subunit interactions for CombFold (instead of AFMv2 in Benchmark 1). The performance on this dataset is comparable to Benchmark 1 (Fig. S2), with Top-1 and Top-5 success rates of 64% and 68%, respectively. In comparison, the Top-1 success rate of AFMv3 is 36%. The fraction of High-quality Top-1 models is higher on this Benchmark (52% vs. 40% for Benchmark 1), indicating that AFMv3 produces pairwise interactions with higher accuracy (Fig. S3), perhaps due to the higher number of recycles and larger training set.

### Integration of experimental data

Integrative structure modeling is often used to determine the structures of large macromolecular assemblies using information from a variety of sources, such as crosslinking mass spectrometry, cryo-Electron Microscopy, or bioinformatics analysis^22, 38–41^. The information is used for scoring and sampling models to produce structures that are consistent with the available data. Here, we add to CombFold support for integrating information about known interactions between subunits and distance restraints that originate from crosslinking mass spectrometry. This type of information can be obtained for individual complexes *in vitro* or for multiple assemblies identified from *in situ* experiments^42–45^. AFM does not currently support the integration of this type of data. Recently, AlphaLink^46^ was developed to add distance restraints support to AlphaFold2/OpenFold as a bias to residue-residue contacts, similar to template support in AlphaFold2. This method requires subsampling of MSA to give more weight to distance restraints and is currently not applicable for complex structure prediction. The advantage of CombFold is that it can integrate additional information during the assembly stage (Methods).

We apply CombFold with distance restraints for human mitochondrial translocase TIM22 (PDB 7CGP), a Benchmark 1 case, for which both CombFold and AFM failed to produce an accurate prediction (TM-score of 0.57 and 0.67, respectively). We used crosslinking mass spectrometry experiment for this complex^47^ to compile a set of 12 distance restraints. We also divided the chains into two groups for assembly (Methods), based on a known structure of a subcomplex of TIM9 and TIM10 (PDB 2BSK). The resulting model is of High-quality with a TM-score of 0.85 (Fig. 2h).

### Accuracy on Benchmark 3 (mostly homomers)

We obtain a Top-1 success rate of 57% on this benchmark, accurately modeling 87 out of 153 complexes (Fig. 3a, Table 1). Moreover, most of the successful predictions (75 out of 87) are of High-quality (TM-score > 0.8). When Top-10 predictions are considered, the success rate is 58% and 82 out of 89 are of High-quality. The higher fraction of complexes with High-quality models compared to heteromeric Benchmarks 1 and 2, demonstrates the challenge of assembling heteromeric complexes with high accuracy where multiple inter-subunit orientations need to be optimized simultaneously. The predicted confidence correlates with the TM-score (Pearson r = 0.44, Fig. 3d). Moreover, the accuracy of CombFold does not decrease with an increase in complex size (Pearson r = -0.09, Fig. 3e).

**Figure 3.**
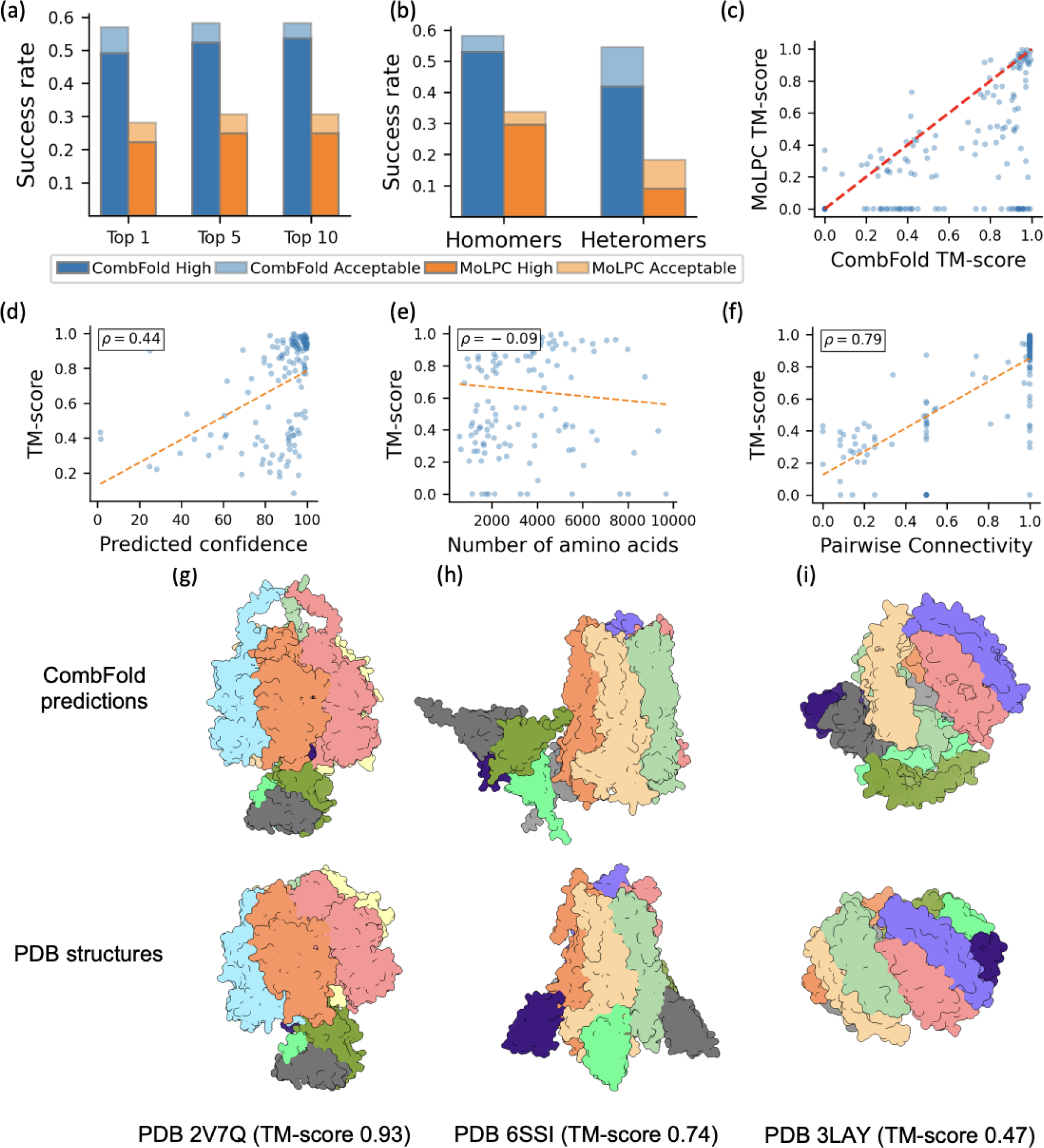
Accuracy of CombFold on Benchmark 3. **(a)** The Top-N (N=1, 5, 10) success rate of CombFold (blue) and MoLPC (orange). **(b)** Top-1 success rate for homomers and heteromers. **(c)** TM-score comparison for CombFold and MoLPC. **(d)** Predicted confidence vs. the TM-score for CombFold. **(e)** The number of complex amino acids vs. the Top-1 TM-score. **(f)** The success rate of AFM in producing pairwise interactions as measured by the pairwise connectivity vs. the TM-score. **(g)** High-quality model of F1-ATPase (top) vs. the x-ray structure (bottom). CombFold prediction contains 159 additional amino acids that are not modeled in the x-ray structure, providing full structural coverage. **(h)** Acceptable-quality model of *Erwinia* ligand-gated ion channel in complex with nanobodies (top) vs. x-ray structure (bottom). The channel is accurately modeled however the location of nanobodies is incorrect. **(i)** Incorrect model of zinc resistance-associated protein from *Salmonella enterica* (top) vs. x-ray structure (bottom).

CombFold success rate correlates with the success of AFM in producing structures of pairwise interactions as measured by the pairwise connectivity (Pearson r = 0.79, Fig. 3f). This correlation is higher than for Benchmark 1 complexes, as in the assembly of homomeric structures, CombFold relies mainly on one or two pairwise interactions. As a result, CombFold accuracy is limited by the reported success rate of ∼60% for AFM in predicting pairwise protein-protein interactions^11–13^. In contrast, in the assembly of heteromeric structures, multiple pairwise interactions are considered, and pairwise interaction can form indirectly even if it is not predicted correctly by AFM. Therefore, the success rate of CombFold on heteromeric complexes is higher (Table 1, Fig. 2).

For comparison, the Top-1 success rate of MoLPC on Benchmark 3 is 28% and Top-10 is 31% (Fig. 3a,c), and the success rate of Multi-LZerD and HADDOCK is 0%^30^. This difference is attributed to our utilization of multiple AFM models and the assembly algorithm that performs a more exhaustive combinatorial and hierarchical search compared to the Monte Carlo Tree Search used by MoLPC. When Benchmark 3 complexes are divided into homomers and heteromers, there is no significant difference for our method, while there is a gap in favor of homomers for MoLPC (Fig. 3b).

### Accuracy on Benchmark 4 (CASP15)

We apply our method to the seven CASP15 targets containing more than 3,000 amino acids. For proteins with a single chain, the chain was divided into domains according to IUPred3^48^, and each domain was used as an input subunit. The method accurately predicted the structure of four complexes using the automatic pipeline (Fig. 4a-d). For target H1111, the N-terminal domain of lcrD protein was removed as it was interfering with the rest of the structure. This domain is not present in the published experimental structure. We find that the pairwise interaction generated by AFM and used by our method to compute structures with cyclic symmetry does not always produce ideal cyclic symmetries (Fig. 4b,c). One future direction is to optimize AFM transformations to ideal symmetries, using symmetric docking tools^49–53^ and selecting docked orientations with a pairwise interaction interface most similar to the one produced by AFM.

**Figure 4.**
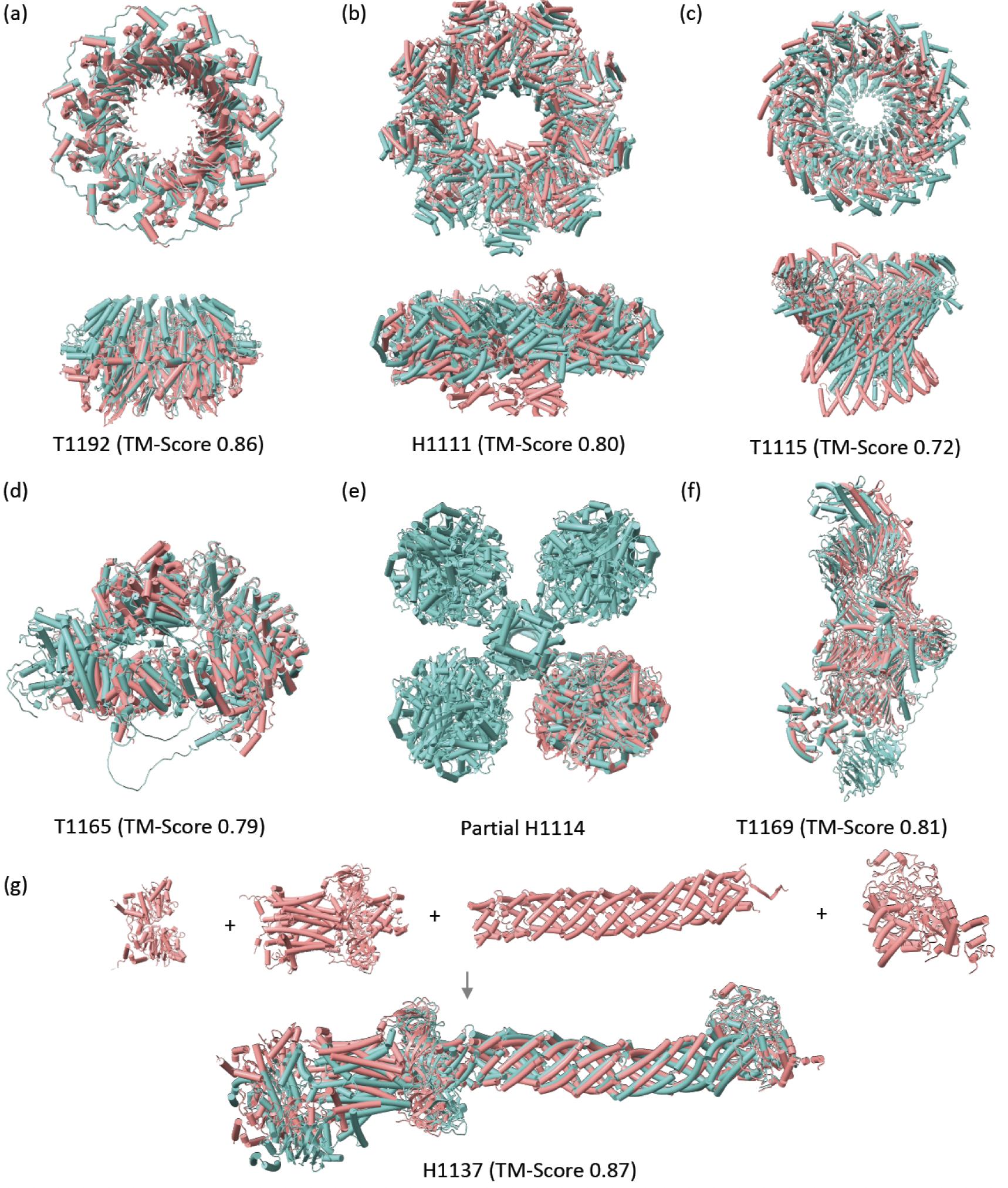
Predicted structure for CASP15 targets. CombFold predictions (coral) are superimposed on the experimental structure (light blue) or top-scoring CASP model if the experimental structure is not released. **(a)** High-quality CombFold model of RAD52 vs. x-ray structure (PDB 5XRZ). **(b)** High-quality CombFold model of YscY-YscX-LcrD vs. x-ray structure (PDB 7QIJ). **(c)** Acceptable-quality CombFold model of stomatin complex vs. top-scoring CASP15 prediction. **(d)** Acceptable-quality CombFold model of Modified ligase Tom1 vs. top-scoring CASP15 prediction. **(e)** High-quality CombFold model of two HucS and two HucL subunits from [NiFe]-hydrogenase Huc complex vs. cryo-EM structure (PDB 7UUS). **(f)** High-quality model of SGS1 vs. cryo-EM structure (PDB 8FJP). **(g)** The four domain groups that were separately modeled by AFM (top) and the final assembly submitted to CASP15 vs. experimental cryo-EM structure (bottom, PDB 8FEE).

The remaining three targets did not have sufficient coverage by pairwise AFM interactions to assemble the complexes. For one of the targets (H1114, PDB 7UUS), a partial subcomplex of two HucS and two HucL subunits could be assembled (Fig. 4e). The other two targets (H1137, T1169) could be predicted with high accuracy by manually dividing their chains into subunits or manually setting subsets of subunits for prediction by AFM (Fig. 4f,g). Here we describe our prediction of the MceG complex (H1137, PDB 8FEE) for which we submitted to CASP15 a single model which had the highest Contact Agreement Score (QS) of 0.90 among all submissions for this target and a TM-score of 0.87. The pairwise AFM predictions enabled us to identify domains and domain-domain interactions, such as helical domains, roughly the same length in six different chains that are folding into a tube-like structure. Interactions between domains that received a low PAE score (<10.0) were used to generate an interaction graph between the domains, enabling us to group the interacting domains into four groups. The domain fragments we used had overlapping amino acids to enable us to connect consecutive domains by structural alignment. In certain cases, the overlap was of dozens of amino acids, and in other cases, it was of entire domains (Table S1). AFM was used to calculate models for each of the four groups, and the overlapping amino acids were used to connect the models of the four groups and obtain the final model (Fig. 4g).

### Application for predicting complexes without known structure

Complex Portal is a database that contains manually curated information on stable macromolecular complexes^31^. We queried the database for all complexes with over 5,000 amino acids, known stoichiometry, and without homology to any experimentally determined structure (Methods) to obtain 28 complexes from three organisms (*Homo sapiens*, *Mus musculus*, *S. cerevisiae*). High-confidence structures were found for seven complexes (Fig. 5, S4).

**Figure 5.**
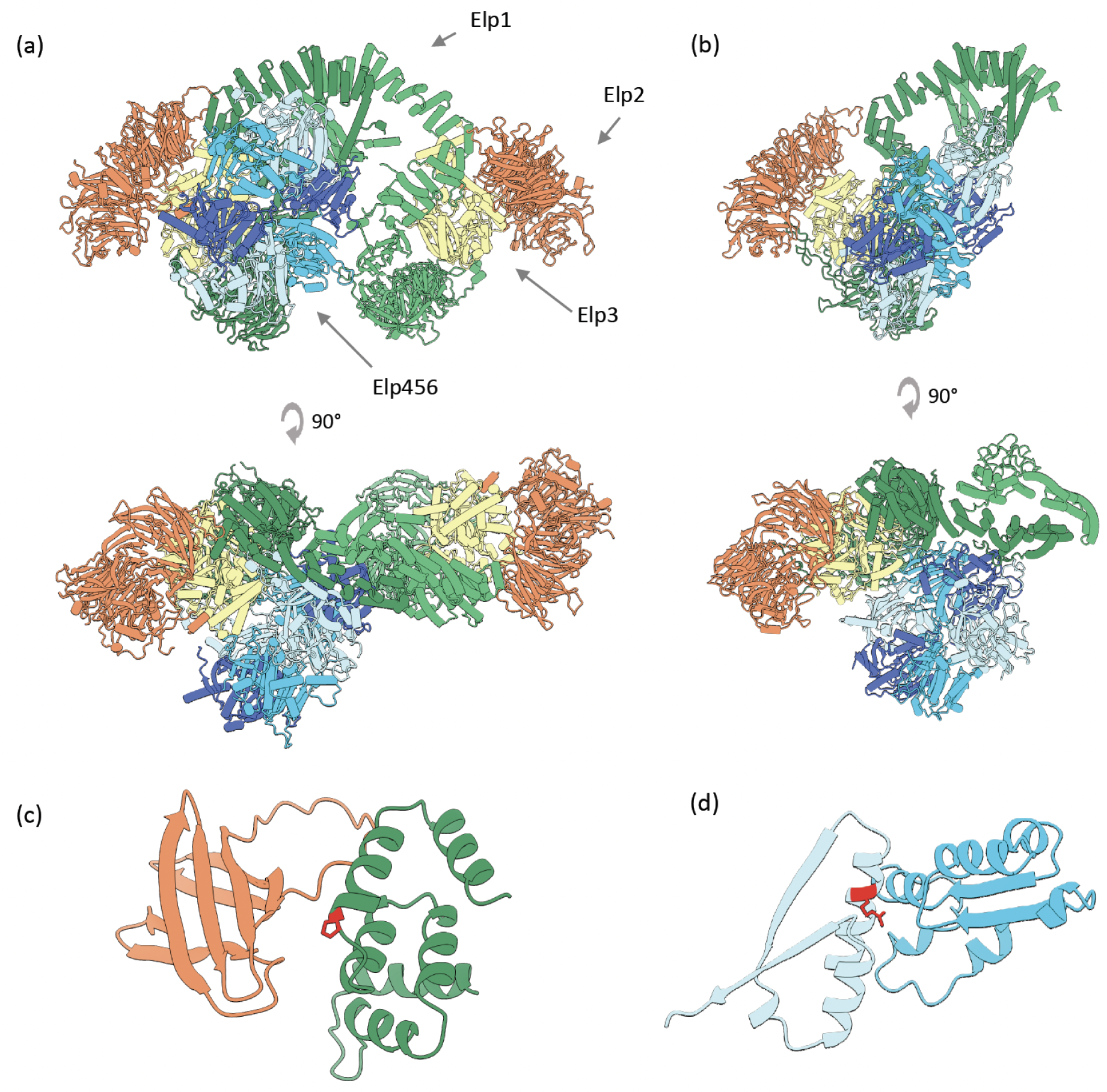
Modeling the human Elongator holoenzyme complex. **(a)** CombFold prediction for the human Elongator holoenzyme complex. **(b)** Part of the complex structure in yeast, as determined by Cryo-EM (PDB 8ASV). **(c)** The interface between Elp1 (green) and Elp2 (orange) with a likely pathogenic mutation P914L in Elp1 is depicted as sticks (red). **(d)** The interface between Elp4 (light blue) and Elp6 (sky blue) with a pathogenic mutation R289W in Elp4 is depicted as sticks (red).

One of the high-confidence predictions is the human Elongator holoenzyme complex which consists of six proteins, Elp1-6, two copies of each. A dimer of Elp123 subunits interacts with the Elp456 subcomplex. Partial homologous structures of *S. cerevisiae* are available, with larger subcomplexes published recently^54^. The structure predicted by CombFold is consistent with the published homologous structure (Fig. 5a,b). Moreover, the predicted structure can be used to explain the effect of mutations. We extracted all the pathogenic mutations from ClinVar^55^ (Table S2) and classified them based on the predicted structure into those that could disrupt protein core or protein-protein interactions (Fig. 5c,d).

### Stoichiometry prediction

The major obstacle to applying our method to known interactions and complexes is the need for stoichiometry information. Our assembly algorithm can be applied to a set of subunits without stoichiometry using the AFM-predicted representative structures and pairwise interactions as follows. Different stoichiometries can be enumerated using the same AFM models as an input and the confidence prediction can be used to estimate the correct stoichiometry. This enables us to perform the resource-intensive AFM calculation once and sample possible stoichiometries with the fast assembly algorithm.

Here we present two examples of this application. The first is the complex of mitochondrial ATP synthase with bound native cardiolipin that contains 10 copies of ATP synthase subunit c forming a symmetrical cylinder (Fig. 6a). We used CombFold to predict complexes with 14 stoichiometries: 2 to 15 copies of subunit c and the correct number of copies for all the other subunits. There is a significant increase in predicted confidence for assemblies with 10, 11, and 14 copies of subunit c (Fig. 6b), indicating that confidence can be used to narrow down the set of possible stoichiometries.

**Figure 6.**
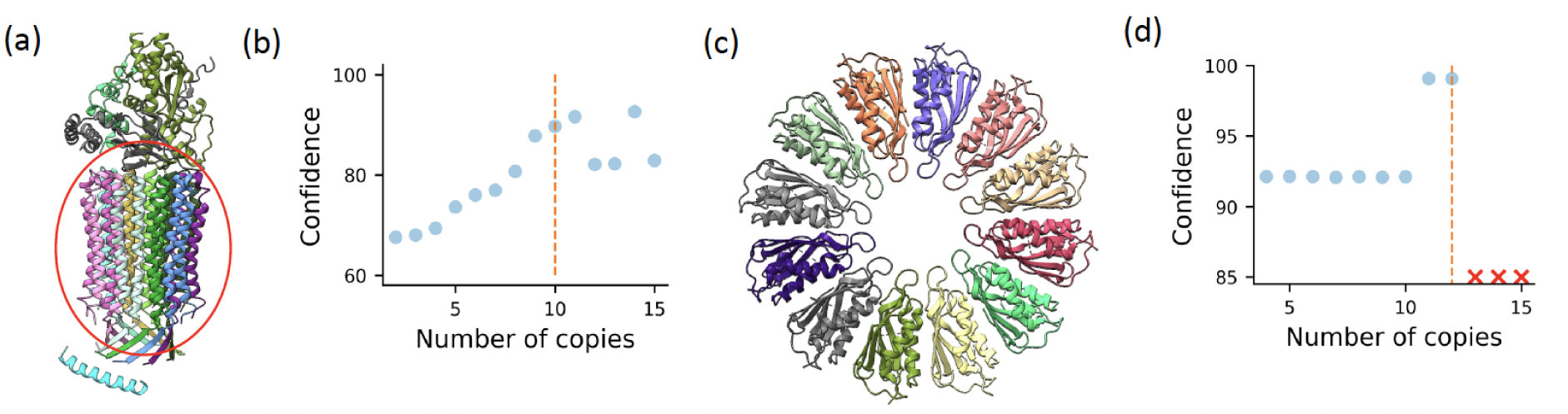
Stoichiometry prediction using CombFold. **(a)** A structure of mitochondrial ATP synthase with bound native cardiolipin (PDB 6TDX). Circled is a symmetrical structure formed from 10 copies of subunit c. **(b)** CombFold predicted confidence as a function of the number of copies of subunit c. **(c)** A structure of PelC dodecamer (PDB 5T11) **(d)** CombFold predicted confidence for PelC dodecamer as a function of the number of copies in input stoichiometry.

Another example is the PelC dodecamer from *P. phytofirmans*. This is a symmetrical complex composed of 12 copies of Lipoprotein (Fig. 6c). We applied CombFold to predict complexes with 14 stoichiometries (2 to 15 copies of the PelC subunit). For 13 or more copies no structure could be assembled without significant steric clashes. There is a spike in the predicted confidence for assemblies with 11 or 12 copies (Fig. 6d). This demonstrates that not only confidence is an indicator of stoichiometry, but that the ability to assemble is another indicator.

## Discussion

We present an approach to predict the structure of large multi-subunit protein complexes based on substructures predicted by AFM for pairs or larger subsets of input subunits. Our method is powered by the combinatorial assembly algorithm that exhaustively enumerates best-scoring assembly trees resulting in accurately predicted assemblies. Moreover, information that can be converted into distance restraints, such as crosslinking mass spectrometry datasets, can be integrated into the assembly algorithm. We validate the approach on four datasets with Top-10 success rate of 57-74% for both homomeric and heteromeric assemblies (Figs. 2-4, Table 1). Moreover, CombFold is able to extend by 20% the structural coverage of experimentally solved large complexes where the modeled structure often does not fully cover the sequences. This enables the application of CombFold to extend the coverage of solved structures.

Most complexes could be assembled by CombFold using single chains as subunits. However, for some complexes, dividing chains into domain-level subunits is beneficial for correct assembly, such as CASP15 targets H1137 and T1169. While our method supports domain-level assembly, the decision of whether to split into domains is left to the user. Subcomplexes are often known based on prior knowledge or can be inferred from single-chain structures, such as intertwined domains in CASP targets H1137 and H1114. In these cases, our method can enforce the specific assembly order to compute the known subcomplexes followed by the generation of the whole assembly.

Currently, our success rate is limited by the ability of AFM to produce pairwise subunit interactions (Fig. 2c, 3f). In this regard approaches that enhance the AFM sampling by enabling dropout at inference can be useful^14, 56^. Additional pairwise orientations might be obtained from pairwise docking methods^25, 57, 58^ as in the original CombDock method^17^. This will enable us to further increase the success rate of our method.

We compare CombFold to other complex structure prediction methods. Docking-based methods such as Multi-LZerD^18^ and HADDOCK^26^ are unable to predict large complexes^30^. When compared to the Monte-Carlo Tree Search assembly (MoLPC) that is mainly applicable to homomeric complexes, our combinatorial algorithm doubles the success rate from ∼30% to ∼60% (Fig. 3a,b). This improvement is particularly significant for heteromeric complexes, where the larger number of subunit combinations leads to an increased number of pairwise interactions. By employing a more exhaustive assembly algorithm, we are able to better enumerate the many possible interactions, resulting in a higher number of accurate assemblies. We also compare CombFold to AFM, which is considered state-of-the-art for predicting entire complexes. We find that AFM is still limited compared to assembly methods by the maximal total length of the complex and lack of diversity in the generated structures. Most complexes that are accurately predicted by AFM, are also accurately assembled by CombFold based on the pairwise interactions from AFM (Fig. 2d).

While some complexes assemble into stable structures, others are dynamic and exist in multiple states. The heterogeneity can be both compositional with subunits that interact transiently or conformational with flexible proteins or a combination of both^59^. Addressing this heterogeneity is still challenging. For example, compositional heterogeneity can be addressed similarly to stoichiometry by enumerating compositions during assembly. The conformational heterogeneity is currently addressed based on additional structural information, such as cryo-electron microscopy^60, 61^, cryo–electron tomography^62^, crosslinking mass spectrometry^63^, and single molecule Förster resonance energy transfer ^64^. Our current implementation can integrate distance-based information into the assembly stage and generate multiple models that are consistent with the data.

Large datasets of experimentally observed protein-protein interactions and assemblies are available from Complex Portal, Corum, and STRING^31, 65, 66^. In addition, crosslinking mass spectrometry is providing large datasets of interactions^67^. These datasets can be used by CombFold, including crosslinks that can be converted into distance restraints and integrated into the assembly stage. While the major bottleneck in applying assembly methods on these datasets is unknown stoichiometry, we demonstrate that our approach can be extended to enumerate stoichiometries (Fig. 6) and we plan to further develop this capability to enable the assembly of complexes without known stoichiometry.

## Materials and Methods

### Dataset preparation

#### Benchmark 1

To prepare the heteromeric dataset, we downloaded all the PDB entries (up to October 2022 - 195,832 entries) and retained only those that contain at least 5 unique (distinct) chains, resulting in 4,997 entries. Next, we removed entries containing DNA or RNA molecules because they can cause conformational changes upon binding to the protein complex, as well as, entries with short (<50 amino acids) or long chains (>1,500 amino acids), resulting in 1,383 entries. We also filtered entries that contained multiple source organisms, such as host-pathogen interactions, as AFM performance for complexes often depends on the availability of paired alignment, leaving 682 entries. To remove redundancy, the complexes were clustered based on sequence identity. An entry was removed if it had a matching entry with an earlier release date and at least 80% of the chains matching with 70% sequence identity, leaving 213 entries. We also removed all entries released before May 2018 because they were part of the AFM training dataset^11^. The remaining 148 entries were further filtered by resolution (<4 Å). Further, we only kept the entries where the numbering and residue type of ATOM records was consistent with the SEQRES records to enable simple and accurate model validation. This filtering protocol resulted in 35 structures that were used as the test dataset for the pipeline (Benchmark 1, Fig. S1a).

#### Benchmark 2

This benchmark was prepared following a protocol used for Benchmark 1 with several changes. First, the PDB entries used are all entries up to March 2023. Second, instead of filtering entries released before May 2018 (AFMv2 cutoff date), we used September 2021 (AFMv3 cutoff date). Also, we did not filter complexes with chains over 1,500 amino acids. This resulted in 25 complexes (Fig. S1b).

#### Benchmark 3

We relied on a dataset by Bryant et al.^30^ that contains 175 complexes. 17 complexes were removed due to incorrectly annotated biological units in the PDB (PDBs 1JYM, 1NLX, 1RVV, 1S3Q, 2E6G, 2EWC, 2VYC, 3HHW, 3KIF, 3R90, 4RSU, 5K2M, 5NFR, 6E7D, 6X04, 6ZEE, 7AJP). In addition, 5 structures of amyloid fibrils were also removed (PDB codes: 6LNI, 6MST, 6OSJ, 6SDZ, 7NRQ) due to the expected compositional and conformational heterogeneity.

#### Complex Portal

The Complex Portal^31^ is a database that curates information from literature on stable, macromolecular complexes. We queried all complexes listed with over 5,000 amino acids with known stoichiometry. Entries with a homologous PDB structure were removed using a stricter threshold for homology to focus on entries that are more likely to be novel. Therefore, we defined a PDB entry as homologous to the Complex Portal entry if at least 66% of the chains were matching with at least 30% sequence identity.

## CombFold method

### Definition of subunits

A subunit is a sequence that can either be an independent chain of the complex or a part of a chain (for example a certain domain). Sometimes it is necessary to divide a chain into a number of subunits - either because the chain is too long to be predicted by AFM or because domains are connected by a long linker and are not in spatial proximity. For Benchmarks 1 and 3 each subunit was a full single chain as defined in the SEQRES segment of the PDB entry. In two cases (PDBs 1I50, 6KWY), the longest chain was too long to be predicted as part of a pair (over 1,500 amino acids) so it was divided into two subunits, one containing the first 1,000 amino acids and the other with the rest. For Benchmark 2, long chains were divided into subunits evenly until every pair could be predicted by AFMv3. This affected five complexes (PDBs 8HIL, 8F50, 8ADL, 8A3T, 7OZN). In the CASP15 benchmark, the chains were divided into subunits according to IUPred3 domains^48^ which were used to identify linkers between domains. The linkers connecting the domains were not included in the prediction. For Complex Portal predictions, the UniProt sequences were divided until each pair of sequences was under 1,800 amino acids.

### AlphaFold2 structure prediction

In the first stage, we run AFM on each possible pairing of the subunits. Proteins, both homomers and heteromers, have the ability to create intertwined structures where the interacting chains exchange small segments or compact protein substructures. These interactions can result in a wide range of quaternary arrangements, including dimers, or higher-order oligomers^68^. To account for this, AFM prediction is applied for larger subsets of three to five subunits as follows. For each subunit, we select the most likely interacting subunits based on the pairwise PAE interaction score and use them to build larger subsets (Methods). Here, we limit our calculations to the total length of input sequences of 1,800 which can be run on standard GPUs.

AlphaFold2 runs were performed using ColabFold^69^ with default parameters (without templates), producing five structures per run. Subunits were inputted as separate chains. For Benchmarks 1 and 3, we used AFMv2 and AlphaFold-ptm to obtain 10 structural models. For comparison to CombFold on Benchmark 1, only end-to-end AFMv2 was used. For Benchmark 2, CASP15, and Complex Portal predictions, we used AFMv3 only, as it was not trained on these targets. CombFold predictions on Benchmark 2 were compared to end-to-end AFMv3.

### Extracting representative subunit structures

Each subunit structure from AFM predictions is ranked based on the mean plDDT score using all predicted structures from AFM runs for pairs and larger subsets. The structure with the maximal score is selected as the “representative subunit structure” for the assembly stage. Additional criteria were examined as possible ranking scores including, the average PAE score for the structure, the maximal plDDT, or the interaction score with other subunits in AFM prediction. There were no significant differences between the described possibilities, the mean plDDT which is easy to calculate and more widely used was chosen.

### Computing pairwise transformations

The method computes for each pair of subunits a list of possible transformations between them based on their interaction models from AlphaFold2. All pairs of subunits are extracted from multi-subunit predictions. For each pair, if it is interacting (Cα-Cα distance < 8Å), the transformation between the subunits is calculated. We can mark the predicted interacting structure for two subunits *A* and *B*, and two representative structures for those subunits *A’* and *B’*. Notice that even though *A* and *A’* are the same molecules, the different interactions in each AFM model will result in different structures and different reference frames for *A* and *A’*. We would like to calculate a transformation between the representatives *B’* to *A’* that will result in the interaction interface as close as possible to that of the examined model pair *A* and *B*. To achieve this, the transformation *T*_1_that aligns *A’* on *A* is calculated by computing the transformation that minimizes RMSD^70, 71^. Similarly, the transformation *T*_2_ that aligns *B’* on *B* is calculated. Finally, the desired transformation is composed as *T*_2_ ◦ *T*_1_^−1^. A problem arises when a subunit has a disordered region - this region will be folded differently in each predicted model, which can affect the alignment and the resulting transformation significantly. Therefore, during the alignment, we consider only amino acids that have a high plDDT score (> 80) or at least half of the amino acids with the highest plDDT.

Each transformation is scored using the PAE score of the two subunits. PAE score is computed by AFM for any two amino acids in the structure, predicting their alignment error relating to each other. The PAE score values are between 0 to 30 with lower values corresponding to a lower predicted error. The transformation score is calculated and normalized to be between 1 to 100 by the equation: *max*{1, 100 − *P*^2^/4} where *P* is the average value of PAE of the two interacting subunits. This expression gives the score quadratic properties so that small differences in low *P* scores (which are usually at least 1) will be meaningful, while for high *P* scores, there is not much difference between the score of transformations as it is predicted to be inaccurate.

Other possibilities for scoring were considered, including using the minimal PAE or the interface PAE of the interacting amino acids only. However, the best results were achieved for the complete average PAE. The average interface plDDT which is widely used^12, 30, 72^ was also a good indicator for RMSD from correct transformation however not as good as the average PAE (correlation factor of 0.59 vs. 0.64, Fig. S5).

## Combinatorial assembly of subunits

The input to the assembly stage is a list of representative structures of subunits and a list of pairwise transformations between subunits. The output is a list of assembled complexes containing all the subunits. If all the subunits can not be assembled, the algorithm outputs partial complexes containing the largest number of input subunits. The assembly algorithm proceeds with *N* iterations, where *N* is the number of input subunits. In each iteration, the size of the subcomplexes created is increased, until the *N*’th iteration, where the subcomplexes computed contain all input subunits.

Each iteration contains three stages: subcomplexes expansion, filtering, and clustering. The first stage creates new subcomplexes based on smaller subcomplexes from previous iterations and pairwise transformations that were provided to the algorithm. Each new subcomplex is scored based on the scores of the pairwise transformations that were used to generate it. The second stage filters assembled subcomplexes with steric clashes between subunits. The third stage clusters subcomplexes with the same subunit composition and saves *K* best-scoring subcomplexes.

### Expansion stage

In this stage, we attempt to connect pairs of subcomplexes that have no overlapping subunits and with the total number of *i* subunits, where *i* is the iteration number. For each pair of subunits in the two subcomplexes (of sizes *k* and *i-k*), a new larger subcomplex is computed for each input pairwise transformation between those subunits. The transformation is applied to all the subunits of the second subcomplex, thus bringing it to the first subcomplex.

There is a special reward for scoring symmetrical subcomplexes with over five identical subunits transformed with the same pairwise subunit transformation. This reward compensates for the assembly being based on pairwise subunit interactions, compared to the full assembly by AFM which is likely to result in lower PAE scores if a symmetrical structure was formed. Therefore, if a symmetric structure was generated based on pairwise subunit transformations, the new score is calculated as (*S* + *S* * (100 − *S*)/100), where *S* is the original score of the transformation.

### Filtering stage

As the pairwise transformations can be at least partially inaccurate, applying some of them can result in subcomplexes with steric clashes or violated distance constraints and restraints. Steric clashes are checked for all backbone atoms with plDDT higher than 80 because the representative structures can contain disordered regions, which are likely to clash with other subunits as they are left static during the assembly. A backbone atom of one subunit is considered as clashing if its center penetrates by more than 1Å into the surface of another subunit. The steric clash test is performed for all pairs of subunits, one from each subcomplex. A subcomplex is filtered if there are over 5% of a subunit’s backbone atoms clashing with another subunit.

Distance constraints are imposed on different subunits from the same chain to enforce sequence connectivity. A subcomplex is discarded if the distance between consecutive amino acids from two subunits is greater than the number of linker amino acids multiplied by 3Å.

### Clustering stage

RMSD clustering is performed to cluster subcomplexes containing the same subunits. We have used iterative clustering, starting from the best-scoring subcomplex with the RMSD threshold of 1Å. However, a default RMSD calculation does not account for multiple copies of the same subunit. This means that for a subcomplex with *p* copies of identical subunits, there will be *p!* equivalent subcomplexes. In this case, to compare the two subcomplexes we need to find the correspondence between copies of subunits from different subcomplexes that minimizes the RMSD. Incorrect correspondence will lead to high RMSD for similar subcomplexes. To avoid the enumeration of *p!* configurations, we implemented a heuristic that superimposes only the centroids of the subunits using starting order subunit correspondence. After the initial superimposition, the correspondence for each pair of identical subunits is swapped and the RMSD is recalculated using centroids. If the RMSD has decreased, we proceed with the new correspondence. The swap process is repeated until there is no further RMSD decrease. The final correspondence between subunits is used to calculate the Cα RMSD between the two subcomplexes.

After clustering, only the *K* best-scored subcomplexes of size *i* will be saved for the next iteration (on the presented benchmarks *K*=100). Clustering aids in diversifying the stored subcomplexes and avoiding the dominance of suboptimal ones in the set of subcomplexes for the next iteration.

### Data integration

To consider known interactions between subunits, we group the input subunits into subcomplexes based on the data. Each such group will be assembled separately, followed by the assembly of the groups and remaining subunits into a larger complex. Therefore, the information is used to enforce a specific assembly order that is consistent with the known interactions.

The crosslinking mass spectrometry information is converted into distance restraints. A restraint is considered as violated if the Cα-Cα distance is above a distance threshold. The threshold is defined by the user based on the length of the crosslinker. A subcomplex is filtered in the filtering stage if it violates some percentage of its restraints (default 70%). In the case of ambiguity of crosslinked residues due to multiple copies of the same subunit, we require that one of the possible distances restrained by the crosslink is below the distance threshold.

## Performance analysis

### Runtimes

CombFold runtime is dominated by the AFM prediction runs for subunit pairs and larger subsets. On Benchmark 1, the average GPU time for AFM predictions was 548 and 1,164 seconds for subunit pairs and larger subsets, respectively, running on NVIDIA A30 with 24GB of memory. However, since our method requires *O(N^2)* AFM predictions for pairs and *O(N)* AFM predictions for larger subsets the average total GPU time per complex was 6,991 and 13,427 seconds for subunit pairs and larger subsets, respectively. It is also important to note that the first stage of CombFold that performs AFM calculations can be trivially distributed into the shorter AFM jobs that can run in parallel. In comparison, the average GPU runtime required for AFM for end-to-end modeling of an entire complex was 4,137 seconds running on the NVIDIA RTX A6000 with 48GB of memory (n=16). It is important to note that the CombFold runtime is higher for heteromeric complexes containing more unique chains compared to homomeric complexes of similar size, as multiple identical copies of a subunit will use the same AFM interaction models. Benchmark 1 is designed to contain heteromeric complexes with many unique chains; homomeric complexes, such as in Benchmark 3, have lower runtimes. For example, a symmetrical structure with 10 identical chains requires much less GPU time in CombFold compared to naive end-to-end AFM (as we only need to run a job for two copies of the chains which is much faster compared to 10 copies). The runtime of the unified representation and combinatorial assembly stages is insignificant compared to the AFM and is on average 80-600 seconds on the different benchmarks on a single CPU. In contrast to the generation of pairwise subunit interactions stage, the assembly stage is faster when for heteromeric complexes with a higher number of unique chains. The assembly time is much faster compared to MoLPC, where the reported average assembly stage takes 13,000 seconds.

### Pairwise connectivity

Given a set of pairwise transformations and a target complex structure, this metric measures how many of the pairwise transformations between subunits from the target complex are present in the set. A graph is built, where each node is a subunit in the target complex and an edge is present if there exists a transformation in the set between those subunits for which the DockQ^73^ score relative to the transformation in the target complex is at an acceptable level (DockQ > 0.23). We calculate the connected components of this graph. The pairwise connectivity ratio is defined as the ratio between the number of amino acids in the largest connected component and the total number of amino acids in the complex. A single connected component in the graph (pairwise connectivity = 1.0) indicates that there are pairwise transformations that can lead to the assembly of the complex. In contrast, multiple connected components indicate that accurate assembly is not possible with available transformations.

### Visualizations

Protein complexes were visualized using ChimeraX^74^. Graphs were created using Matplotlib^75^.

### Data and code availability

CombFold assembly is implemented using C++. The code, Colab notebook, and tutorial for CombFold are available from https://github.com/dina-lab3D/CombFold. Scripts and data for manuscript figures are part of the repository. The PDB codes for Benchmarks 1-3 are also part of the repository.

## Acknowledgments

D.S and B.S are supported by the Israeli Science Foundation (ISF 1466/18). Molecular graphics and analyses performed with UCSF ChimeraX, developed by the Resource for Biocomputing, Visualization, and Informatics at the University of California, San Francisco, with support from National Institutes of Health R01-GM129325 and the Office of Cyber Infrastructure and Computational Biology, National Institute of Allergy and Infectious Diseases.

## Supplementary Materials

**Figure S1.**
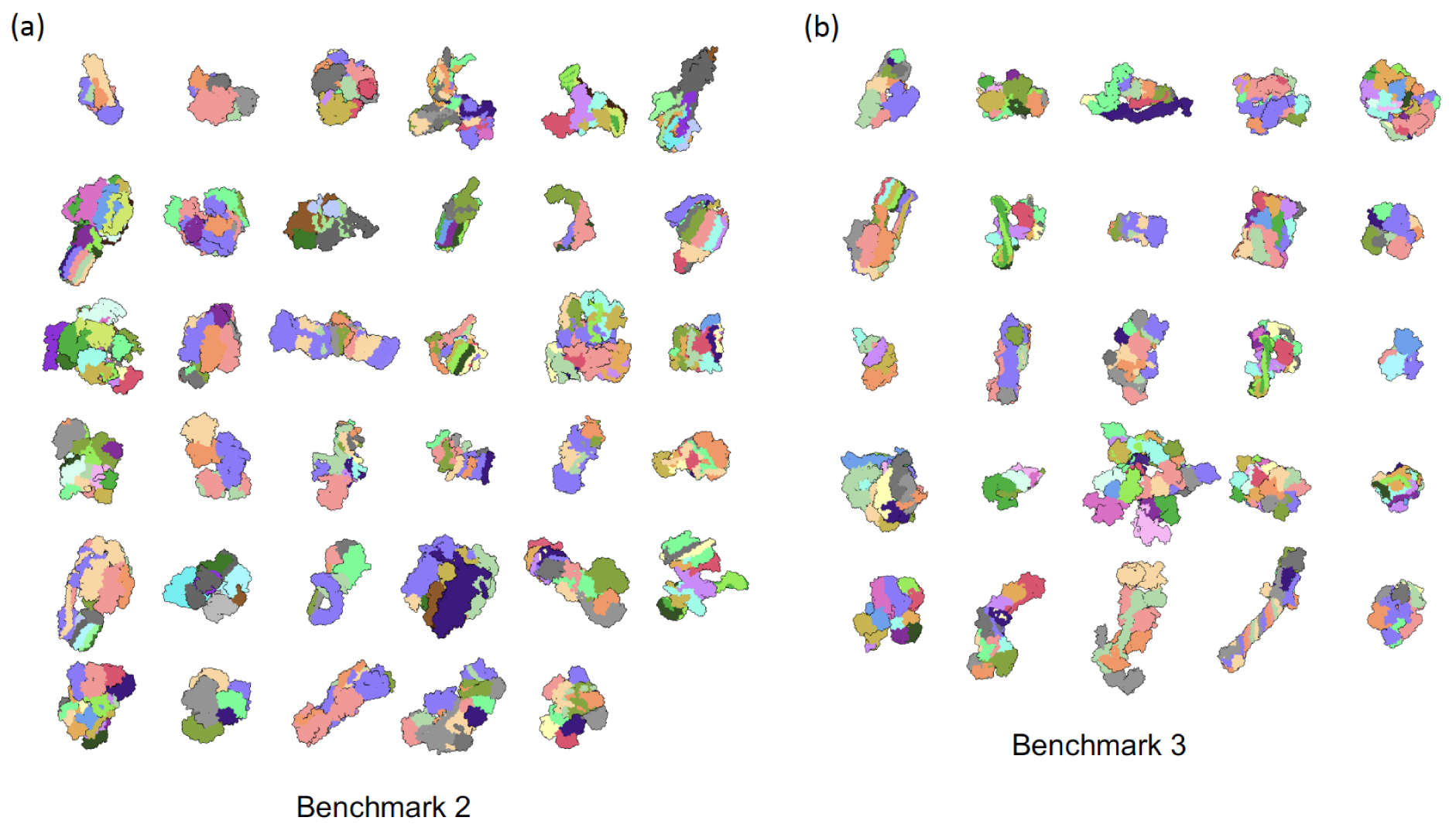
Heteromeric complexes (colored by chain) from **(a)** Benchmark 1 and **(b)** Benchmark 2.

**Figure S2.**
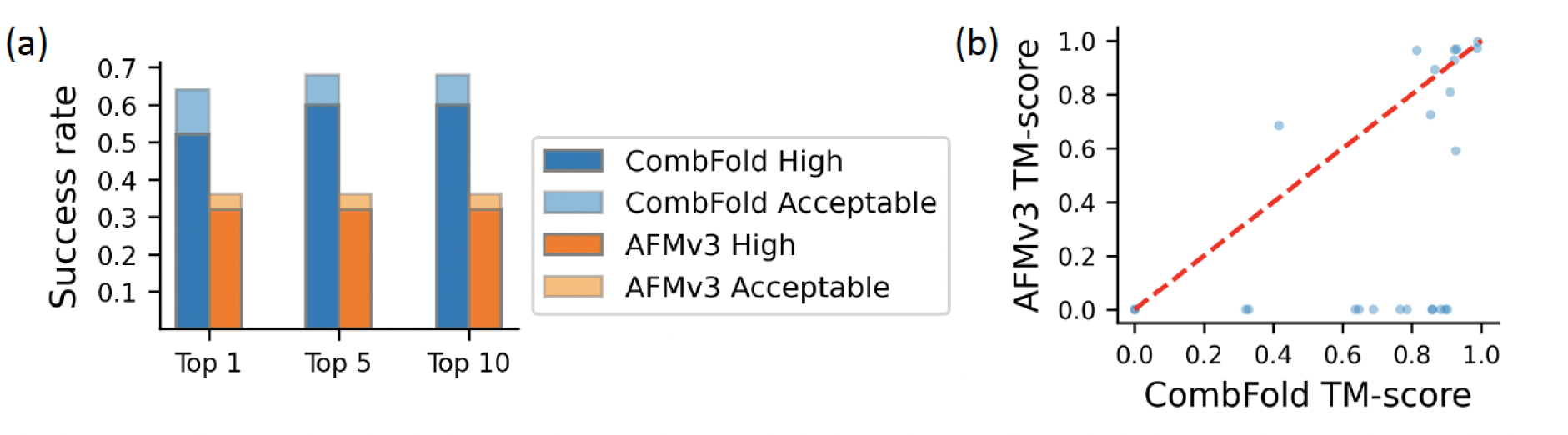
Accuracy of CombFold on Benchmark 2. **(a)** The Top-N (N=1, 5, 10) success rate of CombFold (blue) and AFMv3 (orange). **(b)** TM-score of AFMv3 models vs. CombFold models for Top-5 results.

**Figure S3.**
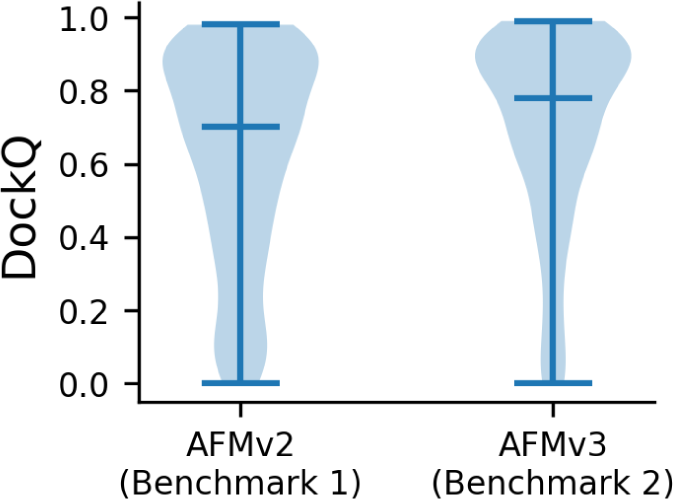
Accuracy of pairwise predictions for AFMv2 and AFMv3. DockQ scores of pairwise interactions predicted by AFM on Benchmark 1 (AFMv2) and Benchmark 2 (AFMv3), for which the PAE-based score is over 50. The median score is 0.70 and 0.78 for AFMv2 and AFMv3, respectively.

**Figure S4.**
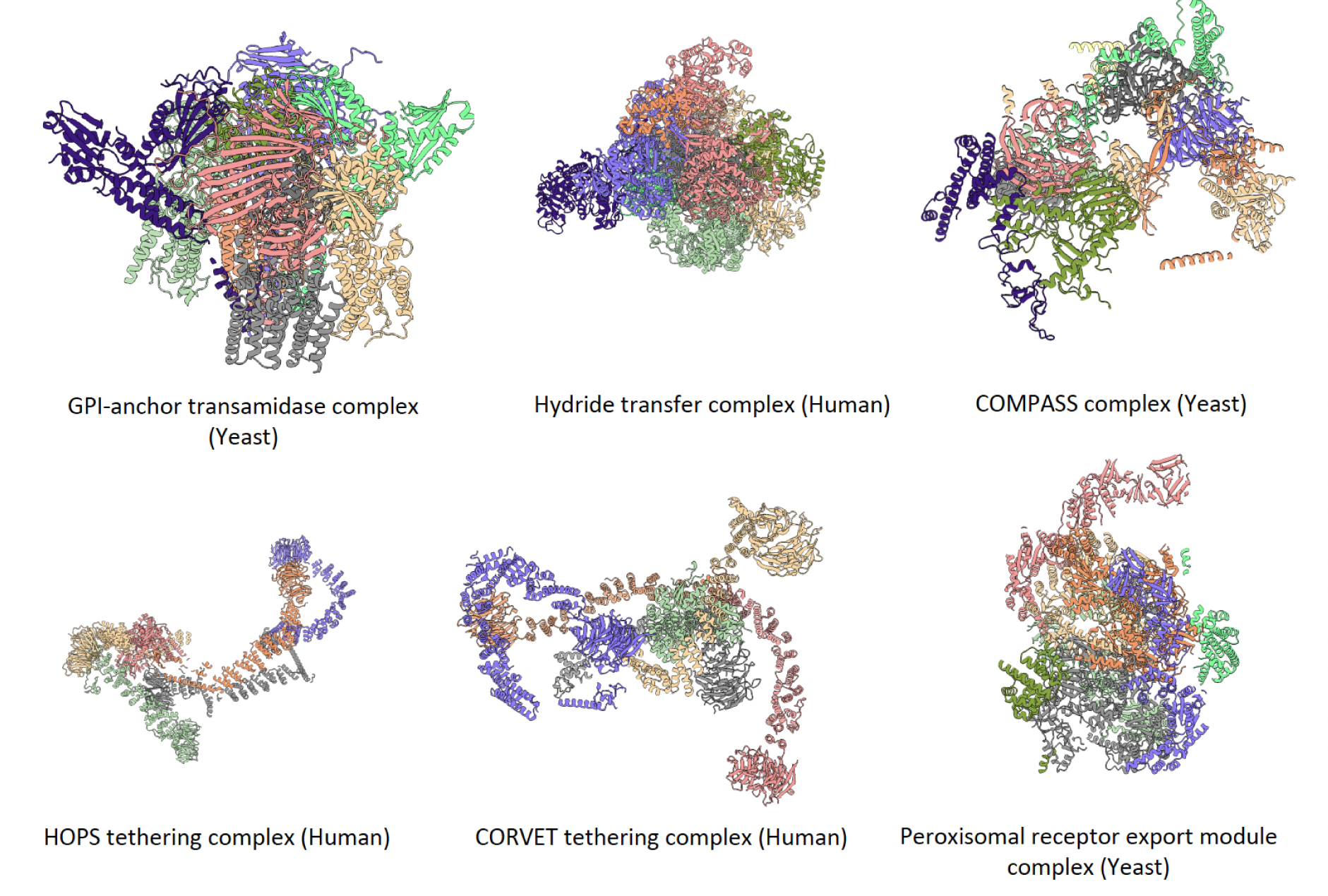
Predicted complexes from Complex Portal with High or Medium confidence.

**Figure S5.**
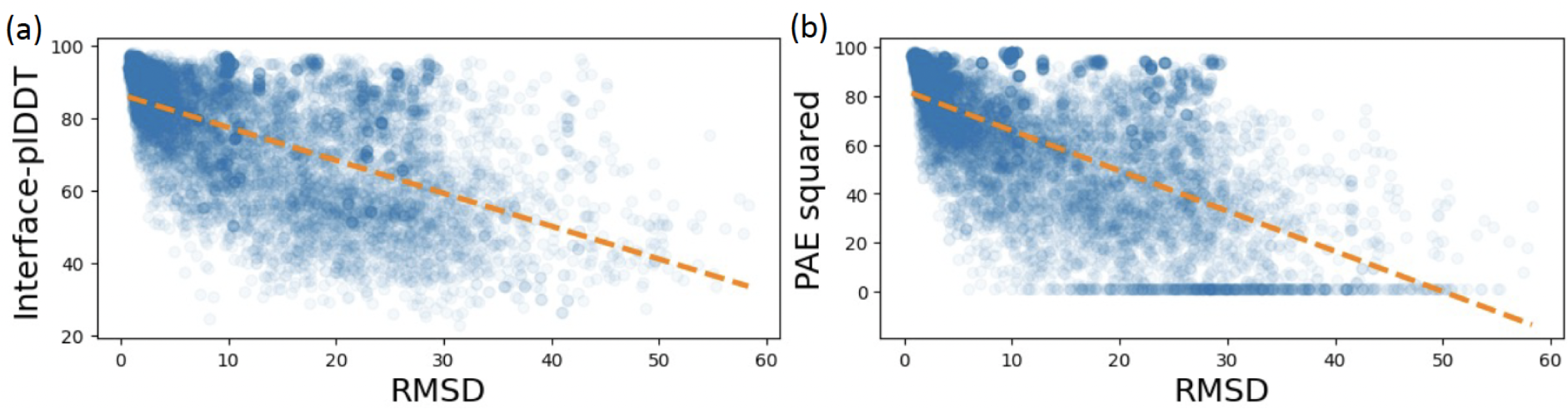
Comparison of average interface plDDT and PAE. (a) Interface-plDDT vs. pairwise RMSD (Pearson r = 0.59). (b) PAE vs. pairwise RMSD (Pearson r = 0.64).

**Table S1:** Subdivision of H1137 into groups of subunits with overlapping domains. Each cell lists the amino acids indexes of the chain present in each model. The division was made so that the helical domain in chains A-F will be calculated as a group (Model 3).

| Chain | Group 1 | Group 2 | Group 3 | Group 4 |
| --- | --- | --- | --- | --- |
| A |  | 1-175 | 155-320 | 300-409 |
| B |  | 1-154 | 134-302 | 282-363 |
| C |  | 1-155 | 135-300 | 280-384 |
| D |  | 1-167 | 147-315 | 295-487 |
| E |  | 1-174 | 154-318 | 298-410 |
| F |  | 1-157 | 137-300 | 280-423 |
| H+I | All | All |  |  |
| G+J | All |  |  |  |

**Table S2:** ClinVar missense mutations for Elongator holoenzyme complex labeled as pathogenic (P) or likely pathogenic (L).

| Gene | Change | Structure position |
| --- | --- | --- |
| (P) Elp1 | Arg696Pro | near interface with Elp6 |
| (L) Elp1 | Pro914Leu | interface with Elp2 |
| (P) Elp2 | His206Arg | interface with Elp1/Elp3 |
| (P) Elp2 | Arg462(Gln/Leu) | core |
| (P) Elp2 | Thr555Pro | core |
| (L) Elp2 | Thr405Ile | core |
| (P) Elp4 | Arg289Trp | interface with Elp6 |
| (L) Elp4 | Tyr91Cys | interface with Elp6 |
| (L) Elp4 | Leu296Ile | core |
| (L) Elp6 | Leu118Trp | near interface with Elp5 |

